# Improved peptide backbone fragmentation is the primary advantage of MS-cleavable crosslinkers

**DOI:** 10.1101/2021.11.23.469675

**Authors:** Lars Kolbowski, Swantje Lenz, Lutz Fischer, Ludwig R Sinn, Francis J O’Reilly, Juri Rappsilber

**Author notes:** These authors contributed equally to this work (Lars Kolbowski, Swantje Lenz).

## Abstract

Proteome-wide crosslinking mass spectrometry studies have coincided with the advent of MS-cleavable crosslinkers that can reveal the individual masses of the two crosslinked peptides. However, recently such studies have also been published with non-cleavable crosslinkers suggesting that MS-cleavability is not essential. We therefore examined in detail the advantages and disadvantages of using the most popular MS-cleavable crosslinker, DSSO. Indeed, DSSO gave rise to signature peptide fragments with a distinct mass difference (doublet) for nearly all identified crosslinked peptides. Surprisingly, we could show that it was not these peptide masses that proved the main advantage of MS-cleavability of the crosslinker, but improved peptide backbone fragmentation that allowed for more confident peptide identification. We also show that the more intricate MS3-based data acquisition approaches lack sensitivity and specificity, causing them to be outperformed by the simpler and faster stepped HCD method. This understanding will guide future developments and applications of proteome-wide crosslinking mass spectrometry.

## Introduction

Crosslinking combined with mass spectrometry (Crosslinking MS) is a powerful tool for detecting protein-protein interactions and the structural characterization of proteins. Many key advances have been made in recent years to expand the complexity of the samples that can be analysed with this technology. These include the database search software^1–3^, FDR estimation^4^ and the enrichment of crosslinked peptides^5–7^. One of the key problems when identifying crosslinked peptides is that one must in principle identify two peptides from the same MS1 signal. The search space is therefore initially very large, comprising every pairwise combination of the peptides that are in the database, i.e. (n^2^+n)/2 crosslinked peptides. This large search space can be reduced experimentally by separating the crosslinked peptides during the measurement by help of an MS-cleavable crosslinker such as disuccinimidyl sulfoxide (DSSO)^8^ or any of its alternatives^9^.

The conceptual advantage of MS-cleavable crosslinkers is evident. The crosslinker readily cleaves upon activation in the mass spectrometer, releasing the individual peptides and thereby enabling the measurement of their individual masses. In the case of the most popular MS-cleavable crosslinker DSSO, the crosslinker cleaves preferentially at two different sites, leading to different crosslinker remnants (also called stubs) for each peptide (**Fig. 1A**). The asymmetric cleavage of this crosslinker produces a pair of alkene (A) and sulfenic acid (S) stub fragments^8^. The S stub fragment commonly loses water forming the unsaturated thiol (T). The two most frequently observed stub peaks per peptide, the A and the T fragment, form a signature doublet signal with a distinct mass difference, allowing their detection and subsequent calculation of peptide masses^10^.

**Figure 1.**
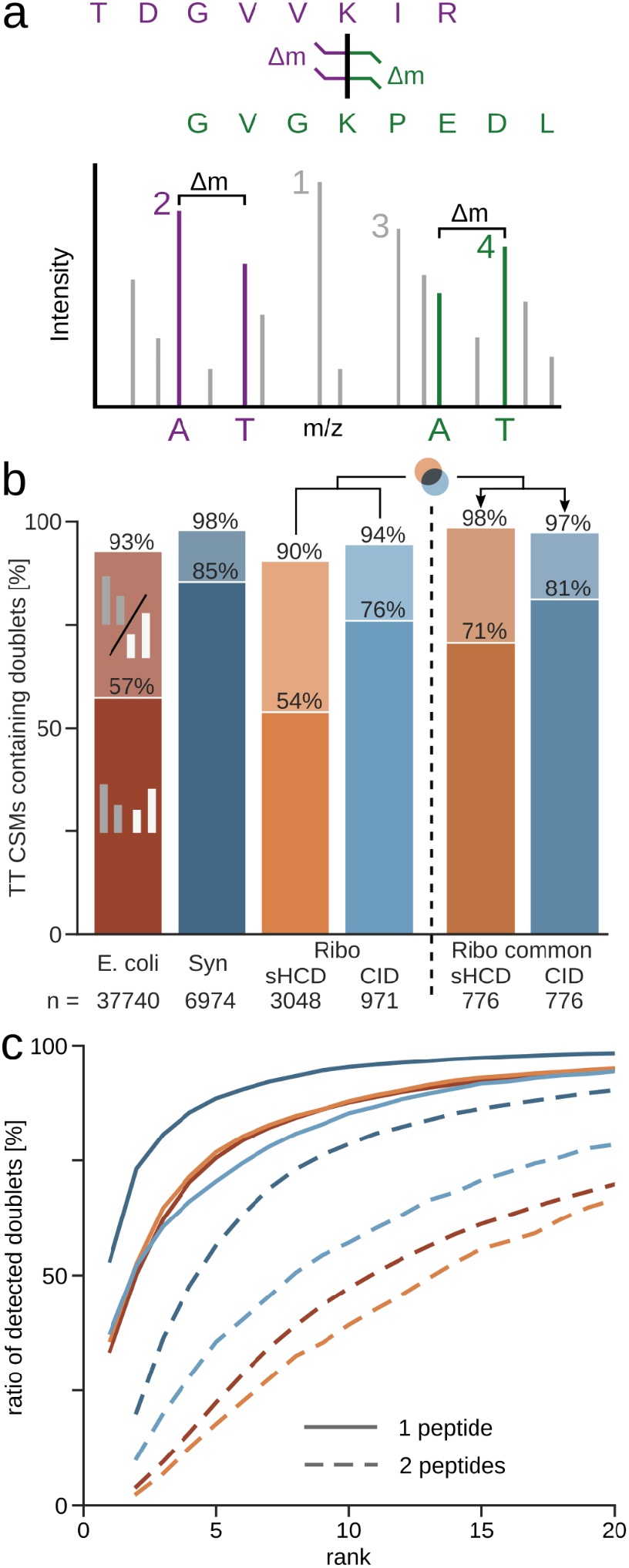
Statistics on frequency and intensity of peptide doublet peaks. (a) Illustration of DSSO cleavage and the resulting signature peptide doublets with the distinct mass difference Δm. Numbers annotate the intensity rank of the peaks, with the rank of the more intense of the doublet peaks being the rank of the whole doublet. (b) Ratio of identified target-target (TT) CSMs that contain one (lighter colour) or both (darker colour) peptide doublets in each dataset (5% CSM-level FDR). Datasets using sHCD are shown in orange-red while CID-MS3 based methods are in blue colours. (c) Percentage of detected doublets passing each intensity rank cut-off. Shown is the cumulative proportion of CSMs containing doublets. Datasets are coloured as in (b). Synapse (Syn); Ribosome (Ribo).

Knowing the individual peptide masses simplifies the database search, as it reduces the search space to pairwise combinations of peptides with these masses. With the individual peptides released in the mass spectrometer, one can also design more intricate data acquisition approaches. The two peptides can be fragmented individually using MS3, which provides separate fragment information of the two - now linear - peptides. For this, generally the crosslinked peptide is fragmented with a low-energy CID fragmentation first, to preferentially cleave the crosslinker instead of the peptide backbone. Then signature doublets are selected for MS3. This approach is routinely employed by studies that use the PIR crosslinker^7^ and DSSO, while some others using DSSO supplement this with a complementary ETD MS2 spectrum^11^.

In an alternative acquisition method, stepped HCD (sHCD), only a single MS2 spectrum is recorded for each crosslinked peptide pair. The peptide is subjected to multiple different collision energies and the fragments are recorded in a single MS2 spectrum. This spectrum should contain the signature doublet (from lower fragmentation energies) as well as additional backbone fragments (from higher fragmentation energies). These spectra can be searched in most crosslinking search tools, with optional filtering for spectra containing cleaved signature peaks during^2^ or after^12^ search.

Despite the clean crosslinker cleavage producing dominant signature peaks in proof-of-concept data of either approach, there is a lack of statistical data of how often this happens in general. It is unclear how many crosslinked peptides give rise to doublets, how prominent these doublets are, and how successful doublet selection is at covering the peptides. It is therefore unknown how many crosslinked spectra are left unidentified when relying on these doublets. sHCD compared favourably to CID methods in the number of crosslinks identified^13^, but a methodical analysis comparing the information contained in their fragmentation spectra is missing and yet is crucial for future design of crosslinkers and acquisition methods.

MS-cleavable crosslinkers have been the tool of choice in many proteome-wide crosslinking MS studies, and it has been suggested that large-scale crosslinking MS depends on MS-cleavable crosslinkers^14^. While conceptually appealing, these advantages and potential limitations of MS-cleavable crosslinkers have yet to be analysed in detail in ‘real world’ scenarios - some comparisons exist, but usually only comparing a few crosslink spectrum matches (CSMs). We systematically investigated the influence of the popular MS-cleavable crosslinker DSSO on the fragmentation of crosslinked peptides. We achieve this by using crosslinker search software that does not rely on the cleaved stubs for identification. This allowed us to clarify how wide-spread the cleavage of DSSO actually is, and to probe the gain of knowing the individual peptide masses for identifying crosslinks.

## Results and Discussion

### Prevalence of peptide doublets in fragmentation spectra of DSSO crosslinked peptides

We analysed three publicly available datasets of DSSO crosslinking experiments coming from three different labs, differing in acquisition method and sample complexity (**Table 1**). The dataset of crosslinked *E. coli* lysate was acquired using sHCD with a low, medium, and high normalized collision energy for each MS2^4^. sHCD is also one of two acquisition methods used to record a dataset of crosslinked, purified 70S ribosomes^13^. In addition to this, Stieger et. al also employed a CID-MS2-HCD-MS3 approach. For this, first a low-energy CID-MS2 was acquired. Then MS3 was triggered when doublets of the correct mass difference (32 Da for A-T) were detected (**Fig. 1A**). Finally, the third dataset called here “Synapse dataset” covered crosslinked mouse synaptosomes and was acquired with a CID-MS2-MS3+ETD-MS2 approach^15^. As in the Ribosome dataset, a low-energy CID-MS2 was acquired for doublet detection. Then, MS3 was acquired as described above, supplemented by an additional ETD-MS2 on the same MS1 precursor.

**Table 1.** Overview of analysed datasets

| sample | crosslinker | acquisition method | variable modifications used in re-analysis | ref |
| --- | --- | --- | --- | --- |
| *E. coli* lysate | DSSO | stepped HCD (sHCD) | oxidation (M), methylation (D,E), deamidation (N, Q), BS3/DSSO -OH; -NH2 (K, nterm) | 4 |
|  | BS3 |  |  |  |
| *M. musculus* synaptosomes | DSSO | CID-MS2-MS3+ETD-MS2 | DSSO -OH; -NH2 (K, nterm) | 15 |
| *E. coli* 70S ribosome | DSSO | stepped HCD (sHCD) | DSSO -OH; -NH2 (K, nterm) | 13 |
|  |  | CID-MS2-HCD-MS3 |  |  |

To assess the prevalence of doublets in the fragmentation spectra of crosslinked peptides, we re-searched the datasets using a search algorithm that does not rely on peptide doublets for crosslink identification. After database search and filtering to 5% heteromeric (inter protein) CSM-level FDR^16^, we looked for signature A and T stub fragment doublet peaks of the identified peptides and the intensity rank of these doublets in each spectrum (**Fig. 1A**).

Even though we did not require doublets to identify crosslinked peptides, they were very common features in our CSMs. We found doublets frequently for at least one peptide, independent of dataset and acquisition method (90 - 98%) (**Fig. 1B**). The CID acquisitions displayed a higher proportion of CSMs with both peptide doublets detected compared to the sHCD datasets. If one looks at only the common identifications of CID and HCD to make up for the difference in number of identifications, the amount of doublets detected for both peptides increases noticeably for sHCD (71%), making the difference to CID (81%) less pronounced (**Fig. 1B**) as does considering only single stub peaks (**Fig. S1**).

We next looked at the intensity of the doublet peaks across these datasets, as this is important for their use during acquisition and data analysis (**Fig. 1C**). In the majority of the spectra, the more abundant doublet is among the most intense peaks, independent of the fragmentation method used. In fact, a doublet peak is frequently the most abundant peak (34 - 53% of the doublet containing spectra). Almost all (94 - 98%) doublet-containing spectra have a peak of the more intense peptide doublet among the 20 most intense peaks.

Spectra typically displayed in publication figures suggest that also the less intense doublet is seen prominently in CID spectra. However, this was only the case for 10% (Ribosome) or 20% (Synapse) of the doublet containing CID spectra of our investigated data. Nevertheless, it is seen among the top 20 peaks in 78% (Ribosome) or 91% (Synapse) of the doublet containing CID spectra. For the sHCD data, the doublet ranks are lower, yet still approximately 70% of spectra have them among the 20 most intense peaks (**Fig. 1C**).

In conclusion, the first doublet is among the most intense peaks for the majority of CSMs independent of the fragmentation method. While the second doublet increases confidence in doublet calling, only one peptide doublet is necessary for deriving both peptide masses given that we know the precursor mass. The visibility of the second peptide doublet is crucial, however, for the successful selection of both peptides for MS3. We therefore investigated how successful selecting doublets from CID-MS2 spectra for MS3 was at covering one or both crosslinked peptides, and if this more complex approach produces more confident identifications than HCD-MS2.

### Speed of HCD outperforms higher sequence coverage of CID+MS3

The ratio of identified doublets and their intensity ranks are important criteria for selecting peptides for MS3 fragmentation. However, absolute numbers of crosslink identifications may also be influenced by other aspects, such as backbone fragmentation and acquisition speed. We used the Ribosome dataset to compare these aspects, as it uses both methods on the same sample. Here, sHCD leads to 1.4 times more residue pairs identified than CID-MS3^17^.

When comparing the common CSMs between CID and sHCD, the overall sequence coverage in sHCD is higher compared to low-energy CID (**Fig. 2A**). This comes as no surprise, as low-energy CID is primarily applied to separate the crosslinked peptides and not for peptide backbone fragmentation. It is intentionally combined with MS3 scans and ETD fragmentation to provide additional sequence information. When we include the corresponding MS3 scans, the sequence coverage increases noticeably compared to that of low-energy CID alone. The overall coverage from combining fragments from CID and MS3 surpasses the sHCD coverage. Therefore, the backbone fragmentation does not explain the higher number of CSMs for sHCD.

**Figure 2.**
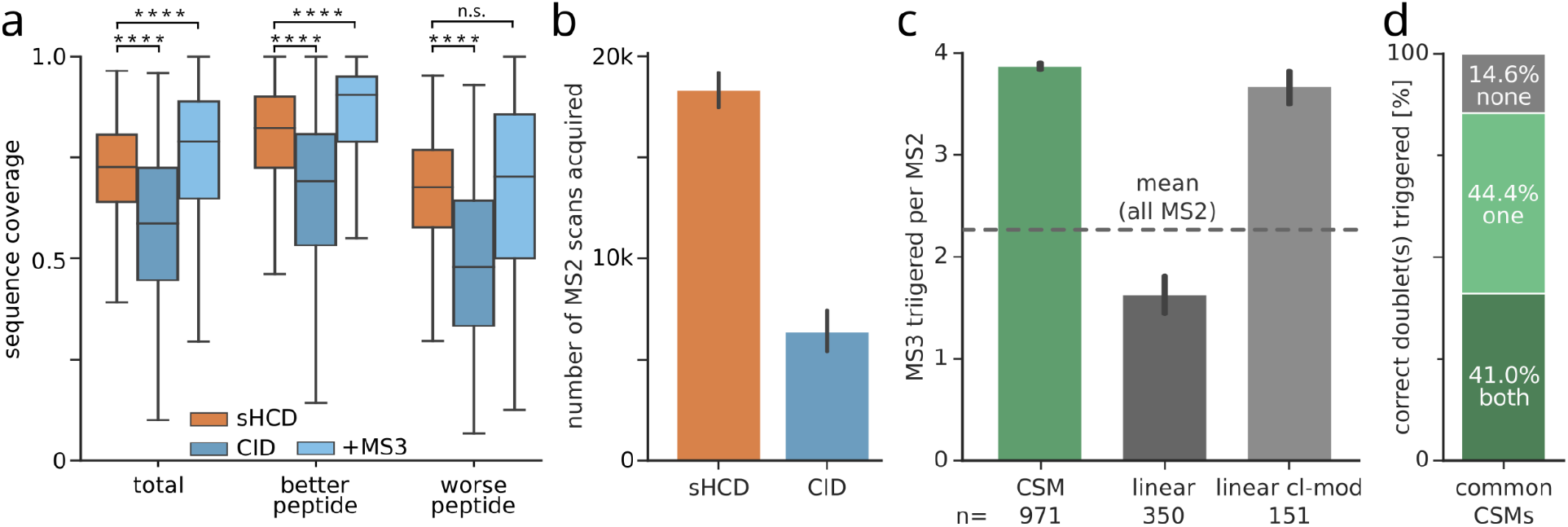
Speed of HCD outperforms higher sequence coverage of CID+MS3 in the Ribosome dataset. (a) Sequence coverage of common CSMs (n=776) identified in both sHCD and CID. Additionally, sequence coverage of CID spectra combined with their respective MS3 scans is shown. Boxplots depict the median (middle line), upper and lower quartiles (boxes), and 1.5 times the interquartile range (whiskers). Asterisks indicate significance calculated by a two-sided Wilcoxon signed-rank (p-value > 0.05: n.s., p-value < 0.0001: ****). (b) Number of acquired MS scans per fragmentation method. Error bars show the 0.95 confidence interval (n=7). (c) Number of triggered MS3 scans per MS2 scan, for CSMs, linear peptide spectrum matches, and crosslinker modified linear peptide spectrum matches, respectively. (d) Proportion of common CID CSMs having no doublets, only one, or both peptide doublets correctly triggered for MS3.

MS3 acquisition schemes require multiple scan and fragmentation events, while sHCD only acquires a single MS2 scan. This difference in complexity and, more importantly, acquisition speed is reflected in the number of total MS2 scans acquired, which on average is almost 3 times lower for the CID-MS3 method, because a lot of acquisition time is spent on acquiring the additional MS3 scans (**Fig. 2B**). The drastically lower sampling of precursors for fragmentation will consequently lead to the reduced detection of crosslinked peptides, which subsequently results in a lower number of crosslink identifications. This is exacerbated by many MS3 spectra being acquired for crosslinker-modified and even for unmodified linear peptides (**Fig. 2C**). Despite this excessive MS3 triggering, for only 41% of the CSMs, MS3 was triggered correctly on both peptide doublets (**Fig. 2D**). This is also reflected in the wider spread of sequence coverage for the worse fragmented peptide, which is crucial for confident identifications of both linked peptides (**Fig. 2A**). Note also that for this peptide the sequence coverage is not significantly increased in CID+MS3 over sHCD.

In this dataset, the speed of sHCD compensates for its slightly lower sequence coverage. sHCD also shows a more symmetric fragmentation of both peptides, as the MS3 approach is limited by its dependency on triggering on the correct doublets. Further development of MS3 approaches should focus on a more sensitive and selective MS3 selection, which in part is governed by the yield of the crosslinker cleavage.

### Peptide doublets for quality control

While some database search algorithms have been built around peptide masses from doublets, others have been built without relying on them. Unarguably, peptide masses are useful information. In an attempt to quantify their value, we investigated the target-decoy CSMs (as representation of the random matches) for the occurrence of peptide doublets. Because heteromeric CSMs are the focus of most biological research questions and are also more challenging to identify, we focused on those for the analysis.

A substantial fraction of random matches have matching peptide doublets (>47% of heteromeric target-decoy CSMs, **Fig. 3A**). However, their extent varies considerably between the datasets. The highest proportion of doublets among target-decoy CSMs is found in the Ribosome dataset (88% or 92% for sHCD and CID, respectively). The *E. coli* dataset contains at least one doublet in 66% of the target-decoy CSMs, while this proportion decreases to 47% for the Synapse dataset. The amount of identified doublets present in target-decoy matches seems less dependent on the acquisition method, and more on the sample and database.

**Figure 3.**
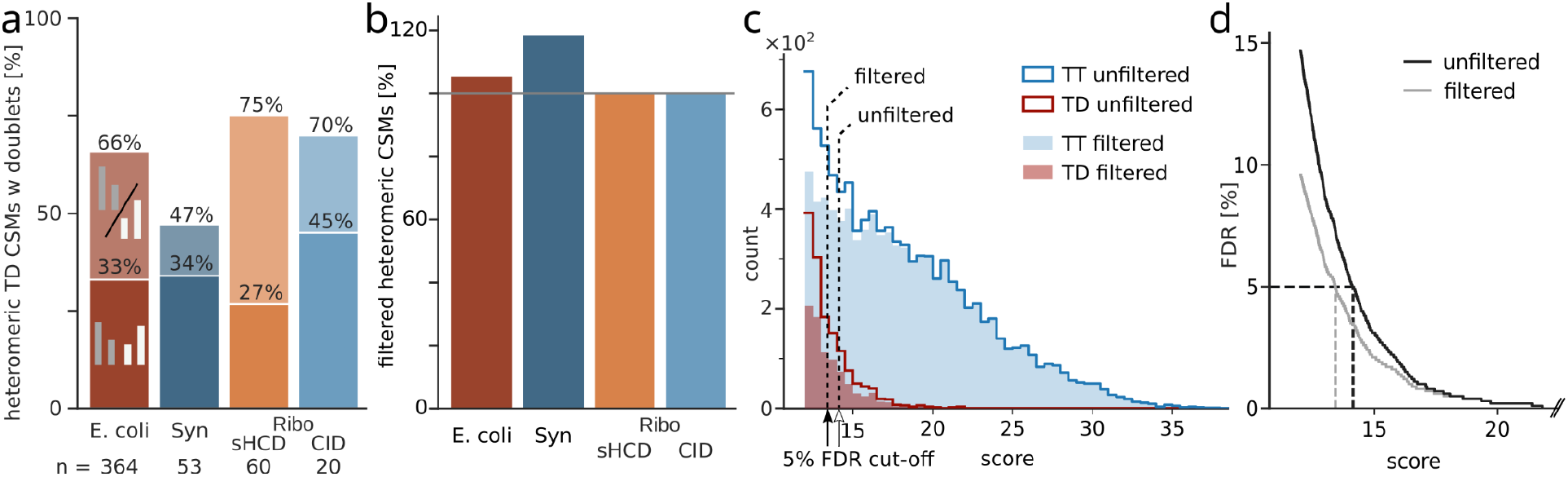
Peptide doublets as a quality control metric for heteromeric identifications. (a) Percentage of heteromeric target-decoy CSMs that contain one or two peptide doublets across datasets at 5% CSM-level FDR. (b) Proportion of heteromeric CSMs at 5% CSM-level FDR when filtering spectra to contain a peptide doublet compared to unfiltered data. (c) Score distribution of heteromeric matches in the *E. coli* dataset. Shown is the distribution of targets and target-decoy matches with and without filtering for peptide doublets. Dashed lines show the resulting score cutoffs at 5% FDR. (d) FDR (interpolated values for visualization) of unfiltered and peptide doublet filtered *E. coli* data. Synapse (Syn); Ribosome (Ribo).

Although heteromeric target-decoy CSMs contain peptide doublets, they do so less often than the heteromeric target-target matches (**Fig. S2**). Based on this difference, we investigated the effect of using this metric as a quality filter. We prefiltered the search results to those spectra that contain at least one peptide with a detected doublet and then re-estimated 5% CSM-level FDR. The gains using this approach are very much dependent on the complexity of the dataset (**Fig. 3B**). Unsurprisingly, the Synapse dataset, which had the least target-decoys containing a matching doublet, shows the largest gains using this approach (19%). However, the *E. coli* dataset only gains 5% in heteromeric CSMs, even though there is a large difference in the proportion of peptide doublets between target and false matches (97% vs. 66%; **Fig. S2, 3A**). This led us to investigate the score distribution of doublet containing matches in more detail (**Fig. 3C**).

The vast majority of high-scoring target-target CSMs contain at least one doublet and are therefore not removed, while targets without a matched peptide doublet tend to have lower scores. In this lower scoring region, there is a steep increase in target-decoy matches, which is only slightly reduced by pre-filtering for a doublet. The effect becomes more apparent when looking at the FDR at different score thresholds. While the increase in error is not as steep for the filtered matches as for the unfiltered, it still grows exponentially (**Fig. 3D**). This holds true also for the Ribosome datasets and to a lesser extent for the Synapse dataset (**Fig. S3-5**).

The moderate gains of using doublets for post-search filtering also suggests that using them during search will offer only moderate gains. Presumably, spectra of high quality, which contain doublets, also tend to contain sufficient peptide fragment peaks so that identification is possible without relying on peptide mass information.

### Comparison of a cleavable to a non-cleavable crosslinker

Non-cleavable crosslinkers are widely believed to be unsuitable for complex samples^14,18,19^. This bases on the assumption that not knowing the individual peptide masses before the search results in the need for exhaustive combination of all peptides in the database and thus an explosion of the search space. However, there are multiple large-scale studies that have successfully employed a non-cleavable crosslinker despite these assumptions^4,12,20,21^. These are based on a detailed understanding of how crosslinked peptides fragment^22^, that offered a computational solution to knowing the individual peptide masses which was then implemented in the search algorithm xiSEARCH^3^. In light of successful usages of both types of crosslinkers, we decided to compare their spectral information to understand any costs and benefits. In addition to DSSO, the published *E. coli* dataset also contains data from the non-cleavable crosslinker BS3. As the data for both crosslinkers were prepared and acquired in a very comparable manner, this dataset offers an opportunity to directly compare the effects of BS3 to DSSO on a complex mixture analysis. Importantly, because of its size and the high number of CSMs identified, the dataset is well suited for statistical evaluation.

A manual side-by-side comparison of CSMs identified in both datasets suggests DSSO to have richer spectra with more fragments. Especially, fragments containing the crosslinking site appear to be more present, mostly as fragments containing an A/S/T stub of DSSO (**Fig. 4A**). We then performed a statistical evaluation of this observation over common CSMs of the two crosslinkers (**Fig. 4B**). This confirmed that DSSO led indeed to a significantly higher sequence coverage than BS3. While the coverage of linear fragments is very similar between the two crosslinkers, the coverage of link site-containing fragments is significantly higher for DSSO. Link site-containing fragments contain the full second peptide (+P) or, additionally for cleavable crosslinkers, just a cleaved crosslinker stub. Indeed, A/S/T stub fragments are the major source of link site-containing fragments for DSSO, while +P coverage is lower than that of BS3. This means that the increased sequence coverage for DSSO stems exclusively from cleaved crosslinker fragments. Crosslinker cleavage appears to promote the cleavage of peptide backbone sites and/or their detection

The better sequence coverage of DSSO-linked peptides improves the separation of true from false CSMs (**Fig. 4C**). For heteromeric matches, DSSO has a larger area under the curve, and especially more high scoring targets, effectively leading to an increase in heteromeric CSMs. While for BS3 3308 heteromeric CSMs were identified, the DSSO dataset resulted in more than twice as many (7316, +121%) (**Fig. 4D**). For self-CSMs, only 29% more CSMs were identified with DSSO than with BS3 (**Fig. S6**), indicating that self-CSMs are approaching exhaustive coverage at the given experimental detection limit. Similar results were seen when including retention time data of heteromeric and self-CSMs^20^.

**Figure 4.**
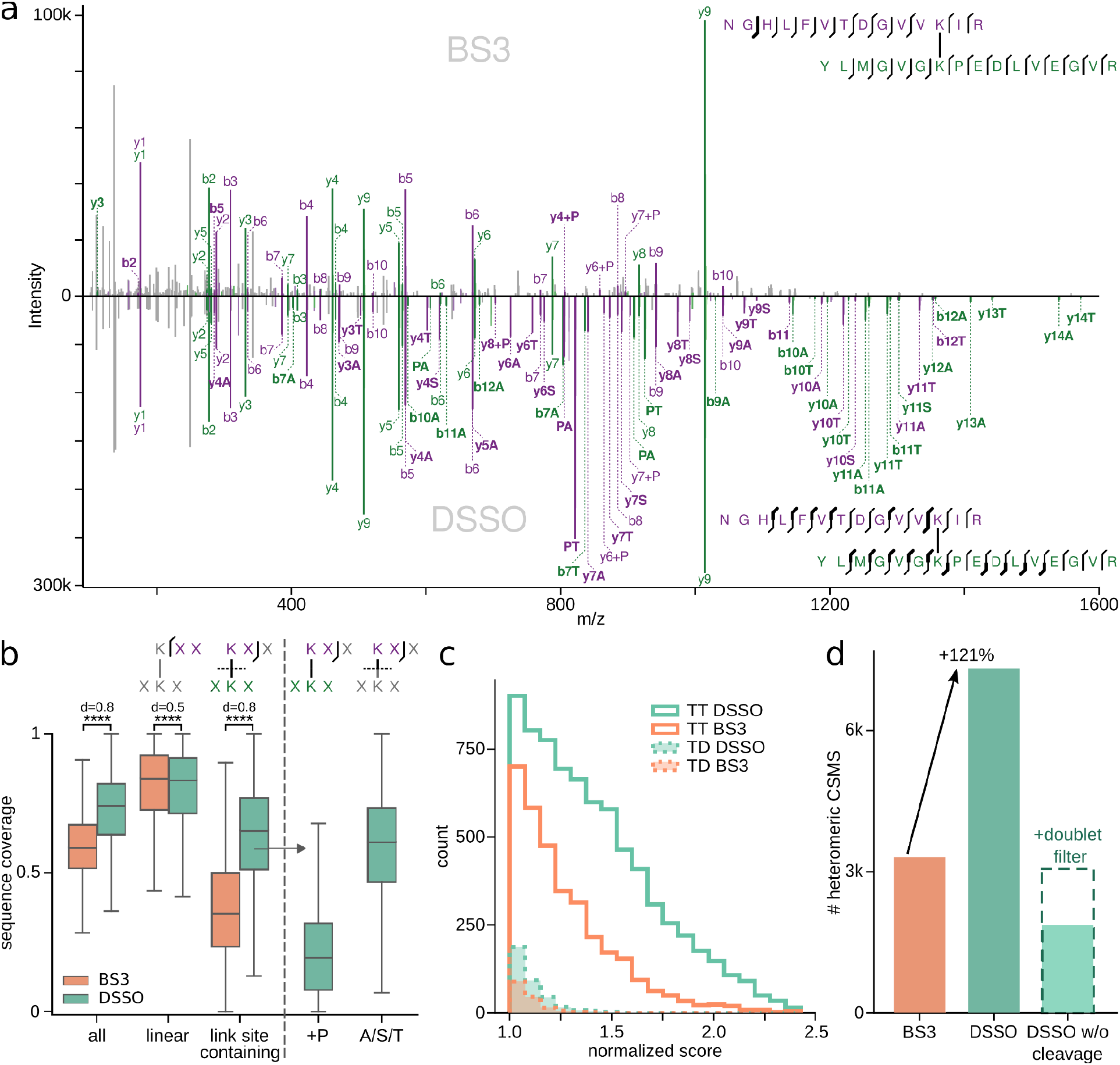
Comparison of non-cleavable crosslinker BS3 to the MS-cleavable crosslinker DSSO. (a) Example MS2 spectrum of a high scoring CSM identified in both datasets. Upper panel shows the CSM from the BS3 dataset. Lower panel shows the same peptide m/z-species identified in the DSSO dataset. Unique fragments are highlighted in bold. (b) Sequence coverage of all, linear and link site-containing fragments (CSMs: n=1437). For DSSO, link site-containing fragments are additionally separated into fragments containing the full second peptide (+P) or only the cleaved crosslinker stub (A/S/T). Boxplots depict the median (middle line), upper and lower quartiles (boxes), and 1.5 times the interquartile range (whiskers). Asterisks indicate significance calculated by a two-sided Wilcoxon signed-rank (p-value < 0.0001: ****). (c) Target-target and target-decoy score distributions of heteromeric CSMs for BS3 and DSSO. Scores were normalized to their respective score cut-off at 10% FDR. (d) Number of heteromeric CSMs passing 5% CSM-level FDR for BS3 and DSSO. As a control, DSSO was additionally searched as a non-cleavable crosslinker and also filtered for the presence of peptide doublets.

To investigate the effect of the cleaved crosslinker fragments on the overall crosslink search performance, we performed another search in which the DSSO crosslinker was treated as non-cleavable. In this search, only 1866 heteromeric CSMs were identified (−74%). Filtering these results for doublet containing results, as described before, increased identifications to 3064. This is, however, still a loss of 58% of CSMs compared to the search considering DSSO as cleavable. Collectively, these observations demonstrate that A/S/T stub fragments play a central role in the success of DSSO for crosslinking mass spectrometry, especially for more complex samples.

## Conclusions

Our work finds a surprisingly limited value of doublet information stemming from crosslinker cleavage for the identification of crosslinks. Nonetheless, we find cleavable crosslinkers to lead to the identification of substantially more heteromeric CSMs. We pinpoint improved sequence coverage as the major contributor to this. This has implications for how to conduct crosslinking studies and the future development of the methodology. Firstly, as many suspected but possibly not for the right reasons, cleavable crosslinkers are preferable for crosslink mixture analyses. Secondly, sHCD is the recommended acquisition method as it achieves almost the same sequence coverage as CID-MS3, but is much faster. CID-MS3 currently lacks speed, specificity and sensitivity. Consequently, future developments of crosslinkers and acquisition methods should focus primarily on sequence information, without compromising acquisition speed. Current choices governing acquisition schemes rely on experimental comparisons, to which we add a methodological understanding of the key parameters that govern crosslink identification. With this, we hope to pave the way for simplified, cost-effective, and standardised workflows that a wider number of labs can use.

## Acknowledgements

The work was funded by the Deutsche Forschungsgemeinschaft (DFG, German Research Foundation) under Germany’s Excellence Strategy – EXC 2008 – 390540038 – UniSysCat and grant no. 426290502 and by the Wellcome Trust through a Senior Research Fellowship to JR (103139). The Wellcome Centre for Cell Biology is supported by core funding from the Wellcome Trust (203149).

## Contributions

S.L., L.K., L.S., F.J.O., and J.R. designed the experiments. S.L., L.K., and L.F. processed Crosslinking MS data; S.L., L.K., F.J.O., and J.R. prepared figures and wrote the manuscript with input from all authors.

## Methods

### Datasets

### Database search and FDR filtering

Mass spectrometry raw data were processed using MSconvert^23^ (v3.0.11729) to convert to mgf-file format. A linear peptide search was employed to determine median precursor and fragment mass errors. Peak list files were then re-calibrated to account for mass shifts during measurement prior to analysis using xiSEARCH^3^ 1.7.6.1 with the following settings: MS1 error tolerances of 3 ppm; MS2 error tolerance of 5 ppm for the *E. coli* lysate dataset and 15 ppm for the others; up to two missing precursor isotope peaks; tryptic digestion specificity with up to two missed cleavages; modifications: carbamidomethylation (Cys, +57.021464 Da) as fixed and oxidation (Met, +15.994915 Da), deamidation (Asn and Gln, +0.984016 Da), methylation (Glu and Asp, +14.015650 Da), amidated crosslinker (Lys and protein N-terminus, DSSO-NH2: +175.03031 Da; BS3-NH2: 155.09463 Da) and hydrolysed crosslinker (Lys and protein N-terminus, DSSO-OH: +176.01433 Da; BS3-OH: +156.07864 Da) as variable modifications; Maximum number of variable modifications per peptide: 1; losses: –CH3SOH, –H2O, –NH3 and additionally masses for crosslinker-containing ions were defined accounting for its cleavability (A: 54.01056 Da, S: 103.99320 Da, T: 85.98264). Crosslink sites for both reagents were allowed for side chains of Lys, Tyr, Ser, Thr and the protein N-terminus. Note that we included a “non-covalent” crosslinker with a mass of zero to flag spectra potentially arising from gas-phase associated peptides^24^. These spectra were removed prior to false-discovery-rate (FDR) estimation. Results were filtered prior to FDR to matches having a minimum of three matched fragments per peptide, a delta score of > 15% of the match score and a peptide length of at least six amino acids. Additionally, identifications of peptide sequences that are found in two or more proteins were removed. FDR was estimated using xiFDR^16^ (v2.1.2) on a unique CSM level to 5% grouped by self- and heteromeric matches.

### Data evaluation

CSMs passing FDR were re-annotated with pyXiAnnotator (https://github.com/Rappsilber-Laboratory/pyXiAnnotator/) with peptide, b-, and y-type ions using MS2 tolerances as described above. The resulting matched fragments were used to check for the occurrence of DSSO A-T doublets and to calculate fragment sequence coverages. We calculated the sequence coverage for our CSMs conservatively, as the ratio of matched N-terminal and C-terminal sequence fragments to the number of theoretically possible sequence fragments (i.e. 100% sequence coverage would mean the detection of at least one fragment from the N-terminal and one from the C-terminal series between all amino acid residues of a peptide). To evaluate the MS3 triggering behaviour the MS3 precursor m/z was extracted from the scan header and compared with the fragment annotation result of the corresponding MS2 CSM. If the MS3 precursor matched a crosslinked peptide stub fragment with 20 ppm error tolerance it was counted as correctly triggered.

