## Supplementary Materials for "Improved peptide backbone fragmentation is the primary advantage of MS-cleavable crosslinkers"

**This file includes:**

Supplementary Figures 1-6

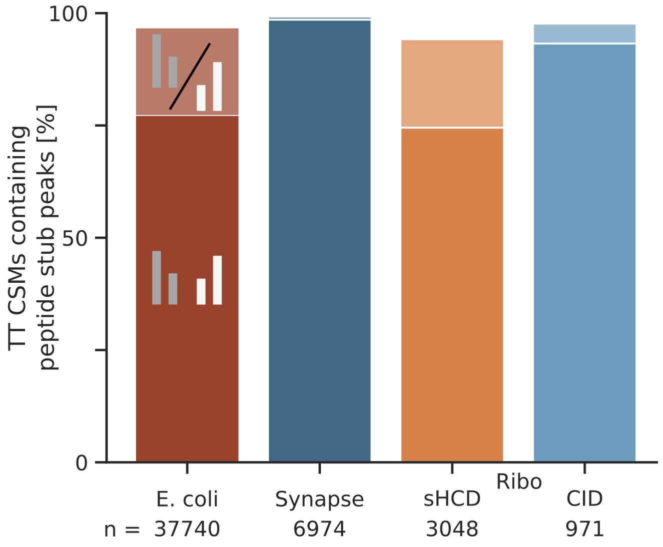


**Figure S1.** Ratio of identified target-target (TT) CSMs that contain at least one peptide stub peak for one (lighter colour) or both (darker colour) crosslinked peptides across datasets (5% CSM-level FDR).

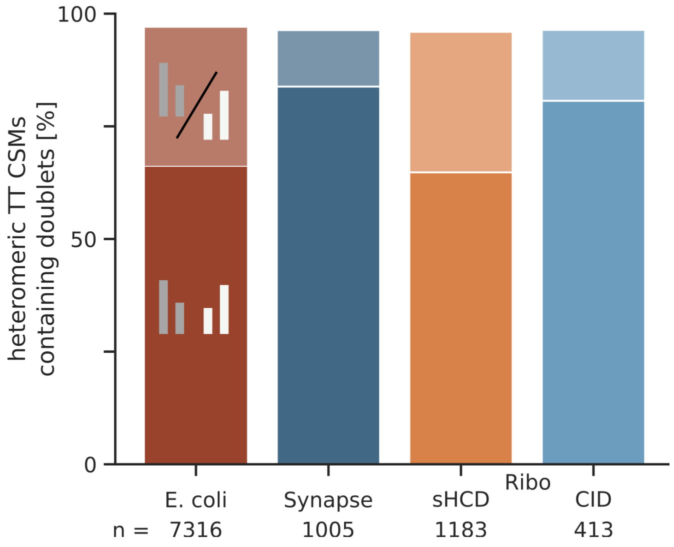


**Figures S2.** Ratio of identified heteromeric target-target (TT) CSMs that contain one (lighter colour) or both (darker colour) peptide doublets across datasets (5% CSM-level FDR).

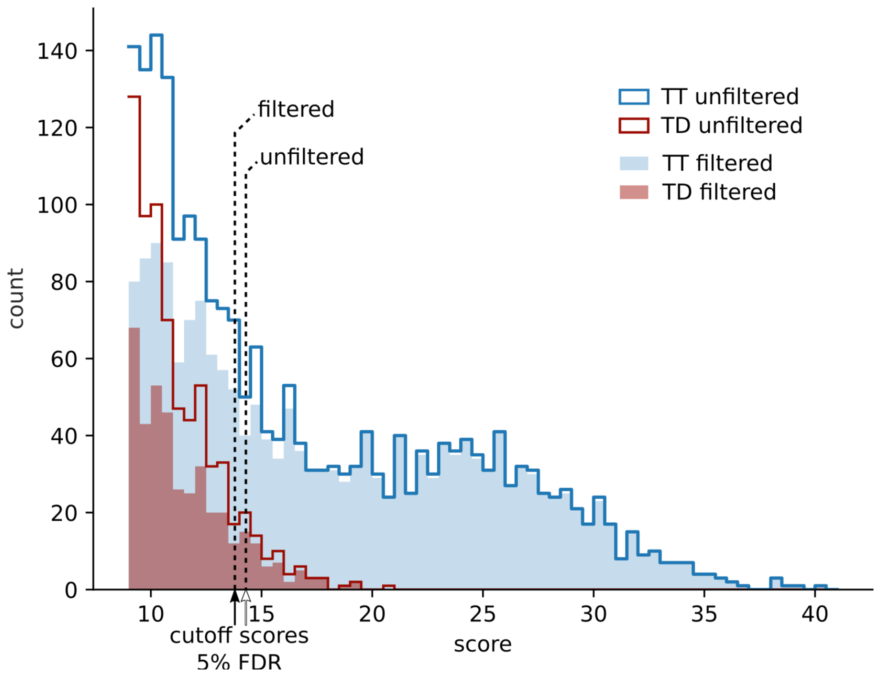


**Figure S3.** Score distribution of heteromeric matches in the Ribosome HCD dataset. Shown is the distribution of targets and target-decoy matches with and without filtering for peptide doublets. Arrows show the resulting score cutoffs at 5% FDR.

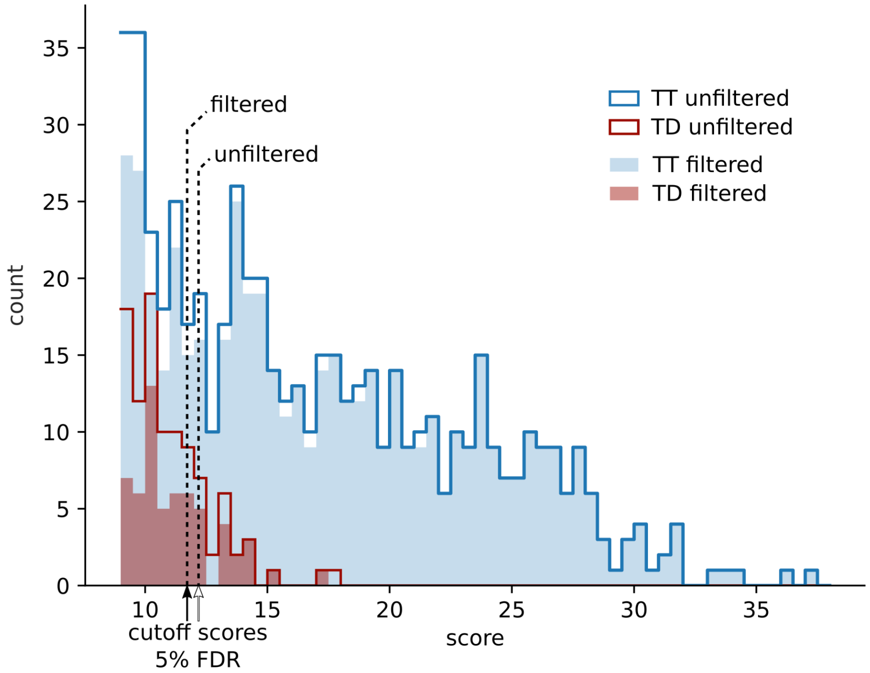


**Figure S4.** Score distribution of heteromeric matches in the Ribosome CID dataset. Shown is the distribution of targets and target-decoy matches with and without filtering for peptide doublets. Arrows show the resulting score cutoffs at 5% FDR.

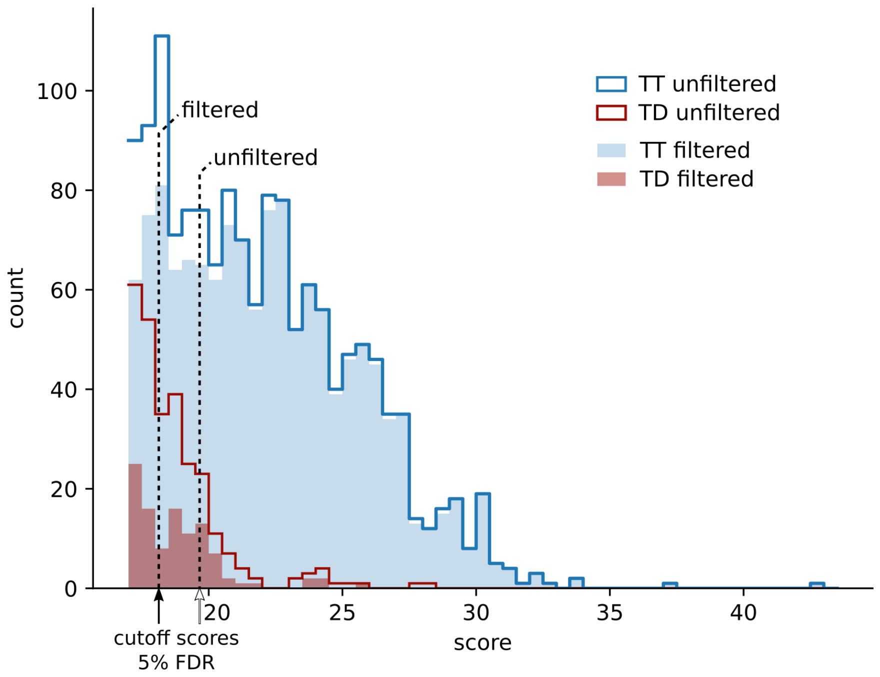


**Figure S5.** Score distribution of heteromeric matches in the Synapse dataset. Shown is the distribution of targets and target-decoy matches with and without filtering for peptide doublets. Arrows show the resulting score cutoffs at 5% FDR.

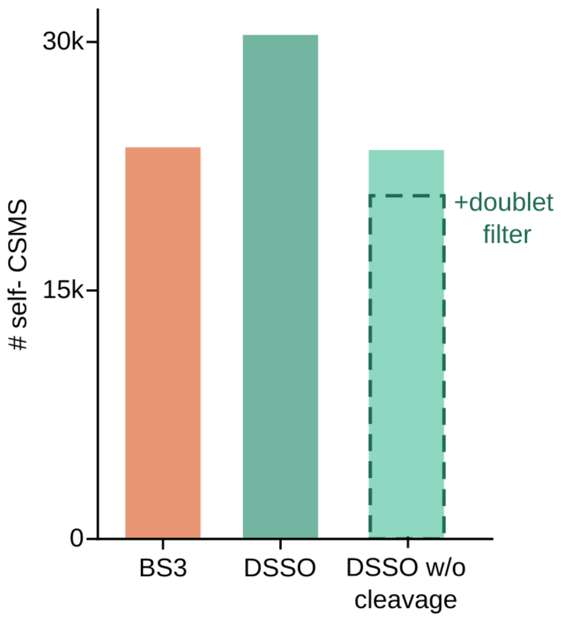


**Figure S6.** Number of self-CSMs passing 5% CSM-level FDR for BS3 and DSSO. DSSO was additionally searched as a non-cleavable crosslinker and filtered for the presence of peptide doublets.
